# Active Strategies for Multisensory Conflict Suppression in the Virtual Hand Illusion

**DOI:** 10.1101/2020.07.08.191304

**Authors:** Pablo Lanillos, Sae Franklin, Antonella Maselli, David W. Franklin

## Abstract

The perception of our body in space is flexible and manipulable. The predictive brain hypothesis explains this malleability as a consequence of the interplay between incoming sensory information and our body expectations. However, given the interaction between perception and action, we might also expect that actions would arise due to prediction errors, especially in conflicting situations. Here we describe a computational model, based on the free-energy principle, that forecasts involuntary movements in sensorimotor conflicts. We experimentally confirm those predictions in humans using a virtual reality rubber-hand illusion. Participants generated movements (forces) towards the virtual hand, regardless of its location with respect to the real arm, with little to no forces produced when the virtual hand overlaid their physical hand. The congruency of our model predictions and human observations indicates that the brain-body is generating actions to reduce the prediction error between the expected arm location and the new visual arm. This observed unconscious mechanism is an empirical validation of the perception-action duality in body adaptation to uncertain situations and evidence of the active component of predictive processing.

## Introduction

Self-body perception is a malleable complex construct, crucial to our interaction with the environment, that emerges from the integration of multisensory stimuli that the brain receives from the physical body^1–3^. The different sensory channels continuously stream information about the state (physiological) and the configuration (postural and in relation to the environment) of the body, and this information needs to be processed and integrated online in order to assure a successful interaction with the dynamic environment in which we live and act^4^. Experimentally induced bodily illusions provide a compelling demonstration of such flexibility, showing how it is possible to transiently induce aberrant bodily experiences—distorted body proportion and impossible postures^5,6^, illusory movements^7^, and the embodiment of external objects^8^ —by manipulating ad-hoc the spatiotemporal correlations of bodily stimuli from different sensory channels.

Hence, one key approach to the understanding of self-body perception is the investigation of body ownership illusions (BOIs)^8^, where external bodily shaped objects like rubber (or virtual) hands are processed as part of one’s own body. These illusions show indeed how the brain can flexibly accommodate for quite coarse sensory conflicts while preserving the integrity of the body percept. In the classic rubber-hand illusion (RHI) experiment^9^, a rubber hand is placed next to the real hidden hand and the illusion is elicited by stimulating both hands with spatiotemporally congruent trains of visuotactile stroking. Once the illusion is established, visual stimuli from the rubber hand (e.g. strokes from a paintbrush^10^ or unexpected threatening events^11^) are processed as if “coming from” the real hand that keeps streaming proprioceptive and tactile sensations. An important element of body ownership illusions is indeed the visuo-proprioceptive binding that is established between visual stimuli from the fake hand and somatosensory stimuli from the physical one^12,13^. Crucially, the illusion is sustained even if the information about the spatial location of the hand provided by proprioception (real hand) and vision (rubber hand) is conflicting^9,14^; and experimental evidence shows how participants undergoing the illusions recalibrate the proprioceptive and visual encoding of the hand location towards the rubber hand^15^. This effect, known as proprioceptive drift, has been interpreted as a way to “explain away” the sensory conflict and can be neatly accounted for within predicting processing computational frameworks as the result of prediction error minimization^16^.

Effects akin to the proprioceptive drift observed in the classic RHI have been consistently reported across different experimental studies whenever the real and the fake limbs were spatially misaligned^13,17–19^. For example, during a full-body ownership illusion experienced over a virtual body initially collocated with the real one, participants previously instructed to be still showed an illusory perceived change in their posture after seeing their embodied avatar slowing moving into a new posture, even in absence of actual movement^13^. In line with this, in a classic RHI experiment in which participants were explicitly instructed to keep their posture and not move, Asai and colleagues showed how participants undergoing the ownership illusion tend to unintentionally apply a force in the direction of the rubber hand^20^. Unfortunately, the experimental design was not able to fully disambiguate whether the observed forces were indeed active strategies for sensory conflict compensation or the result of body posture corrections due to a body midline effect^15,21,22^, as the only location tested was with the rubber hand placed close to the body midline. Besides, automatic and unconscious motor adjustments to reduce conflict between the vision of the arm and proprioception were also reported^23^. Still, if confirmed this form of automatic compensatory movements can be regarded as an active strategy for suppressing the sensory conflict that may arise in bodily illusions, an alternative to “passive” perceptual recalibrations such as the proprioceptive drift.

Similar forms of active compensatory movements have been also observed during intentional actions in participants controlling the movements of an embodied virtual avatar. A clear example is the “self-avatar follower effect”^19^, a motor behavioral pattern observed in immersive virtual reality when users embody a virtual avatar and control it with a many-to-one visuomotor mapping. In particular, participants could control the extension and bending of the virtual arm, while the virtual shoulder rotation was controlled by a Virtual Reality (VR) system independently on the real shoulder configuration (so that the two arms were not spatially overlapping). Under this condition, when asked to perform a sequence of reaching actions, participants tended to integrate within the correct task execution an additional motor component irrelevant to the task, but functional to bringing the real arm closer or overlapping with the virtual one. This motor component cannot be directly explained by optimal control theories of movement^24,25^ but finds instead a natural interpretation in the active inference framework^26^ as an active strategy for suppressing sensory prediction errors^19^. We hypothesize that the brain resolves the sensory conflicting information about the body posture by exerting involuntary active strategies that physically reduce the conflict following a similar principle as the proprioceptive drift.

The aim of the current study is to provide a theoretical account for the active component of the RHI described above, consisting of an implicit (not intentional) attempt to reduce the visuo-proprioceptive conflict arising from the misalignment between the real and the embodied fake hand. We cast the problem in the normative framework of active inference^27^, implementing a model able to predict the force that an agent will produce at the level of the arm when exposed to a classic RHI configuration, where a spatial mismatch between the real and the fake hand is present and actual movement of the physical hand is “locked”. We further conducted an experimental study based on a customized setup integrating a robotic manipulandum with a VR system programmed to induce an immersive virtual version of the RHI^28^. The setup allowed to systematically measure the force generated using a precise force-torque sensor. Force assessments can be then used for formally testing the hypothesis that the active effects observed during illusory ownership (e.g. the attempt to move the hand towards the embodied misplaced hand) are due to the minimization of the prediction error.

Active inference^29^ (AIF), a mathematical framework inspired by the Predictive (Bayesian) brain^30^ that accounts for both perception and action, has been successfully applied, for instance, to model goal-driven behavior in organisms^31^ and the physiology of dopamine^32^. Moreover, AIF accounts of intentional reaching behavior in humans have been proposed^27^ and investigated on humanoid robots^33,34^. Relevant for this work, it was shown how tuning the relative influence of the sensor modalities affected the motoric response in a hand-target phase matching in the presence of visuo-proprioceptive conflict^35^. Perceptual illusions in humans have been also investigated under this paradigm, such as the force-matching illusion^36^. In a previous study, we synthetically replicated the proprioceptive drift observed in the RHI^16^, supporting the general view that this perceptual recalibration results from the minimization of the prediction error produced by the visuo-proprioceptive spatial mismatch between the real and the rubber hands once the illusion is elicited. Here we extend this formulation to the active inference framework and show how the extended implementation can reproduce the experimentally observed forces applied by the hidden real arm in the direction of the rubber/virtual hand.

## Results

We first sketch the main features of the computational model and approach (Figure 1A) and show the model predictions when simulating the virtual hand illusion (VHI). Model predictions are then compared with observations from a dedicated experiment (Figure 1C) in which the VHI was elicited as in classical paradigms (via visuo-tactile stimulation, VT) and the force applied by the hidden arm during the illusion was measured. Although previous studies have shown that illusory ownership over a virtual hand spatially aligned with the real hand can be triggered also without the need for VT stimulation^12^, we opted for the classical paradigm because we needed the illusion to be sustained also in the presence of spatial conflict. Several studies have indeed shown how in these cases, synchronous VT stimulations trigger significantly stronger illusions with respect to conditions with any tactile stimuli^37,38^. In our study we need to retain a spatial conflict to facilitate the appearance of the proprioceptive drift, which in turn is the trigger for the active compensation strategies under testing. Results from simulations and experimental assessments were compared for different spatial configurations of the real and virtual hand (Figure 1B), adopting different degrees of VT congruency (synchronous vs asynchronous) to control for the role of the ownership illusion. Altogether our results show that our AIF model can account for both passive (perceptual drift) and active (attempt to move the hand towards the embodied misplaced hand) effects observed in the laboratory.

**Figure 1:**
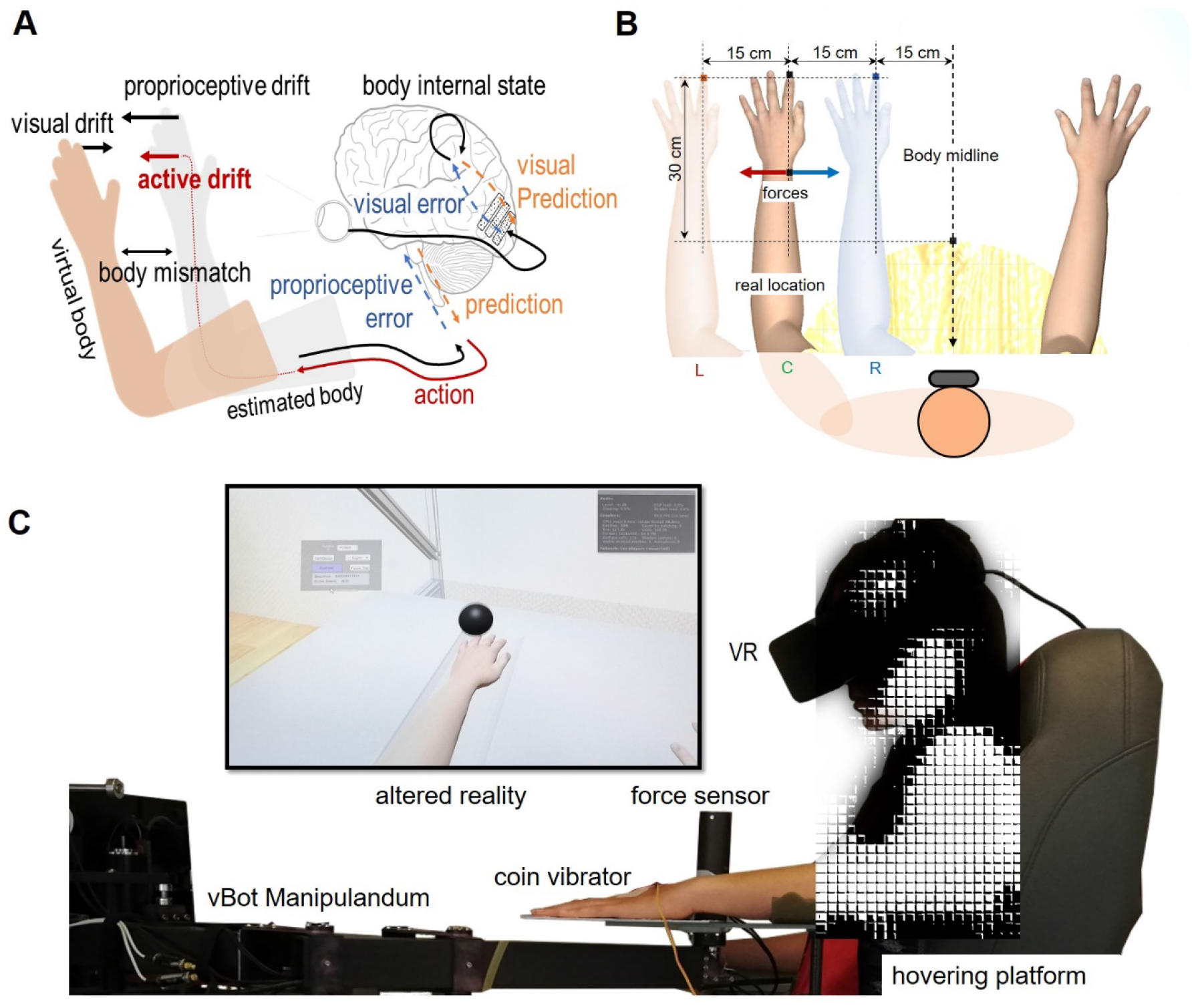
Predictive brain approach and VHI experimental setup. (**A**) Schematic of the perceptual drifts (visual, proprioceptive) and the active component, and its relation to prediction minimization. Both the *proprioceptive and visual drifts* are produced by the same process of body estimation under visuo-proprioceptive mismatch and the *active drift* (forces towards the virtual hand) is a consequence of proprioception error minimization through the reflex arc pathway. (**B**) VHI conditions. At each trial the virtual hand was placed in one of the three different locations: Left (−15cm), Center (0cm) and Right (+15cm) with respect to the real hand location, i.e., the center corresponds to the real hand position. The real hand was placed 30 cm away from the body midline. (**C**) The perceptual stimulation was performed using immersive VR and a coin vibrator for the tactile stimulation. Lateral force measurements were recorded using a 6DOF force-torque sensor on the robotic manipulandum vBot.

**Figure 1:**
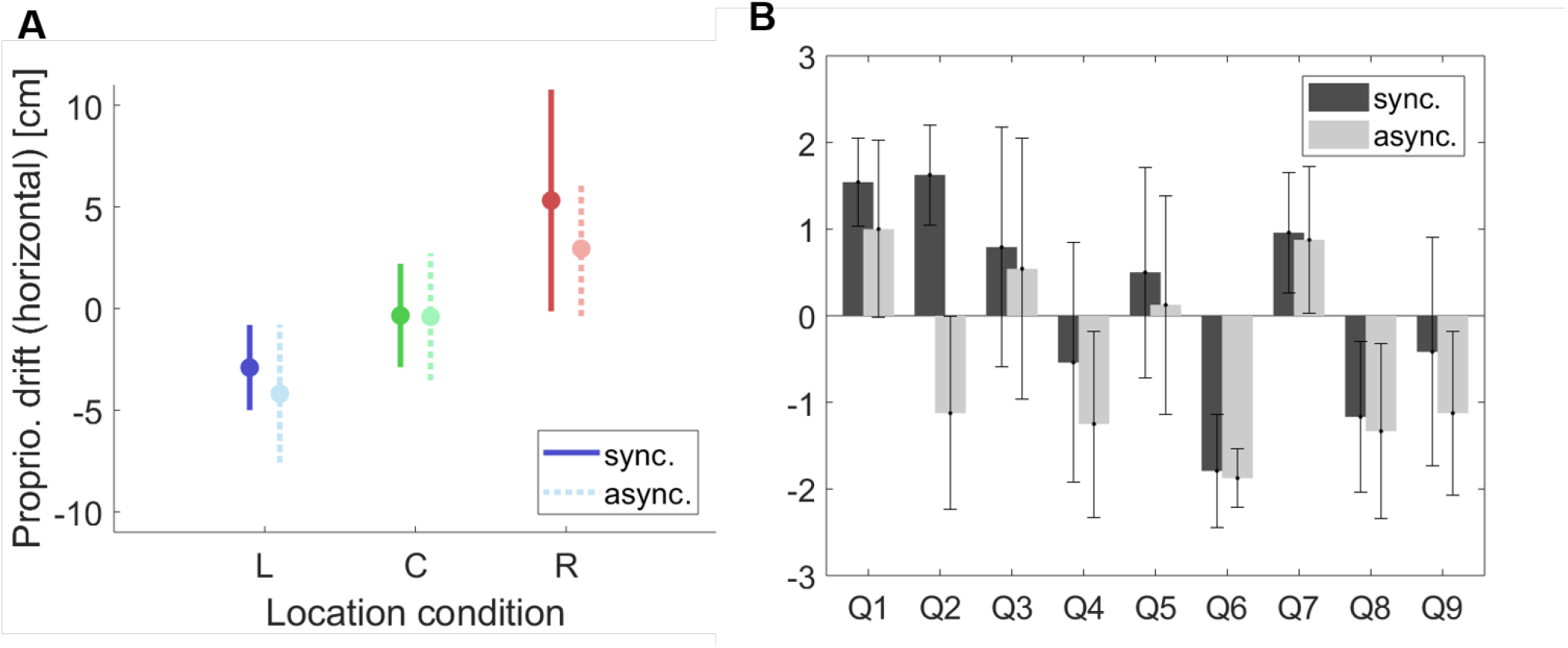
(**A**) Proprioceptive drift for synchronous (continuous line) and asynchronous (dashed line) for the three VR arm location conditions (Left, Center, and Right). (**B**) Body-ownership scoring results from a questionnaire answered by the participants using a 7-point Likert scale (3 indicating strong agreement and −3 indicating strong disagreement).

### Computational model predictions

We model perception and action as an inference process using an AIF^27,39^ based algorithm, sketched in Figure 2A—see the details in Methods. During stimulation, the real hand location is hidden (grey hand) and thus, only proprioceptive cues provide information about the current body pose. The virtual hand is treated as a possible source of information and integrated into the arm posture estimation as a visual cue weighted by causality, which in turn was considered as a proxy for the ownership illusion and modulated as a function of VT synchrony. According to the model, the force at the level of the arm arises as a consequence of the prediction error generated in the multisensory integration process of the visual arm location input and the predicted arm location from the estimated body joint angles.

**Figure 2:**
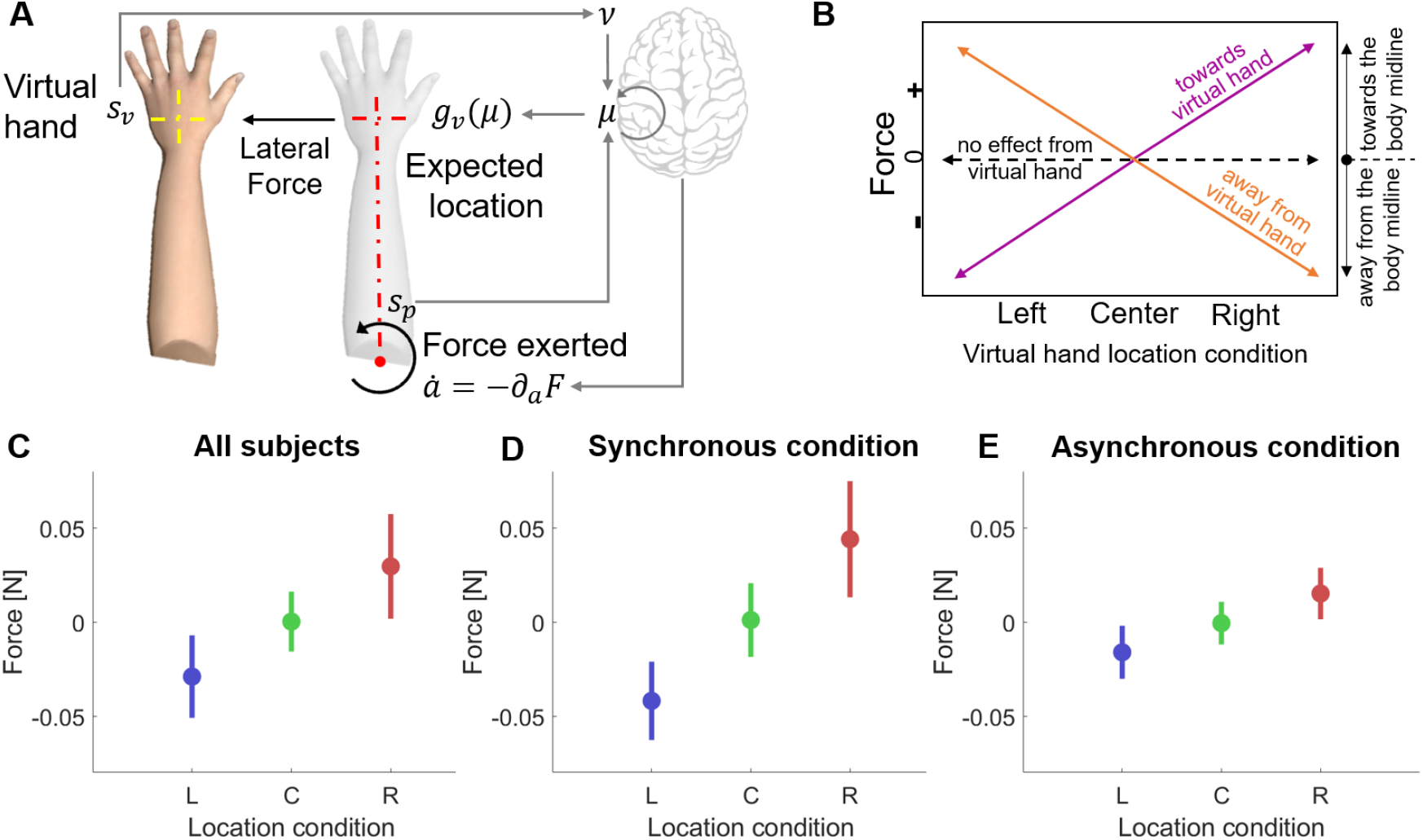
Computational model predictions. (**A**) Arm model description with one degree-of-freedom (DOF) and the VR hand as the visual input. Using the proprioceptive input *s*_*p*_ (joint angle) and the prior information of the body state *p*(*μ*) the model predicts the expected location of the hand *g*_*v*_ (*μ*). However, the visual information (virtual hand, *s*_*v*_) manipulated to appear in a different location produces a sensory conflict: my hand is not at the predicted location. Thus, producing an error that is propagated and generates the perceptual drifts and the active component. According to our hypothesis, the torque direction corresponds to reducing the prediction error: expected hand location minus the VR location. (**B**) Relation between the direction (sign) of the predicted force and the virtual hand location. Our model predicts a trend similar to the one shown by the purple line, e.g., negative force in the left condition implies that the attempted force is towards the virtual hand. (**C**) Mean and standard deviation of the predicted force with the *active inference* model for Left, Center and Right conditions. (**D**) Model predicted mean force for the synchronous condition. (**E**) Model predicted mean force for an ideal asynchronous condition.

The simulated VHI experimental design, had three different visual conditions (described in Figure 1B), corresponding to three different locations of the virtual hand: Left, Right and Center (i.e. in the same location of their real hand). The output of the model is the predicted lateral force, being positive when it is towards the body midline (torso) and negative when it points away from the body. Figure 2B depicts the expected outcome depending on the force sign and the virtual hand location condition. In the left condition, a negative force indicates that the predicted force is towards the virtual hand and a positive one that the hand would move away from the virtual hand. In the right condition, this is inverted: negative force implies that the attempted force would make the real hand go away from the virtual hand and positive force indicates towards the virtual hand. In both left and right conditions zero force indicates no effect of the virtual hand location. Finally, in the center condition positive and negative forces would imply movement away from the virtual hand. In the presence of our proposed active strategies, we should see a trend similar to the one shown by the purple line.

We simulated 20 participants (10 synchronous and 10 asynchronous). Participants’ variability was modeled by assigning different proprioceptive weighting (i.e. the precision or inverse variance of the proprioceptive cue) and randomizing the length of the limb. The results, summarized in Figure 2C-E, predict lateral forces in the direction of the virtual hand in all conditions. Figure 2C shows the mean force for all simulated participants split by the location of the virtual hand (Left in blue, Center in green and Right in red), indicating three differentiated patterns. As expected, in the Center condition, characterized by congruent visuo-proprioceptive information (i.e. no conflict) the model is compatible with no force applied (mean is not significantly different from zero). Conversely, when a spatial conflict is present, the model forecasts an action towards the artificial hand to correct the error of the predicted arm location, which we should observe in human participants. Furthermore, the model in the asynchronous condition predicts attenuated forces (Figure 2D vs Figure 2E), as the asynchronous condition was associated with a suppression of the ownership illusion and therefore a lower level of reciprocal causality between visual and proprioceptive stimuli.

### Active RHI experiment with human participants

To validate the model predictions, we investigated the exerted forces of fourteen human participants in the VHI experiment presented in Figure 1. Participants wore an immersive virtual headset through which they could see a virtual body placed in an anatomically plausible position with respect to their visual perspective. Analogously to the computational model experiment, participants were presented with three different conditions, corresponding to three different locations of the virtual hand (Figure 1B): Left, Right and Center. Center corresponds to the same location of their real hand. Participants were instructed to keep their arm still; the real arm was placed on a locked manipulandum equipped with a 6DOF force-torque sensor (Figure 1C) so that the implicit (non-intentional) horizontal forces applied by participants could be measured (sensitivity: 1/24 Newtons). As we were only interested in the active component of the illusory experience, to obtain enough force sample points for statistical robustness (120 samples per individual), no other typical measures of the RHI such as proprioceptive drift or subjective assessments were taken. However, to complement this analysis, we did a preliminary study with eight participants to assess the perceptual drift and the subjective level of body-ownership from the same experimental setup—See supplementary information.

#### Lateral forces analysis

We first describe the registered lateral forces with respect to time during stimulation averaged for all participants (Figure 3) and second, we present the mean forces for each condition averaged for the stimulation interval 10s-15s (Figure 4). Figure 3 shows the lateral force profile during the stimulation period of 40 seconds, averaged for all participants and trials. Although both synchronous and asynchronous conditions had clear right vs left force patterns, the variability was higher in the asynchronous case producing noisier force profiles.

**Figure 3:**
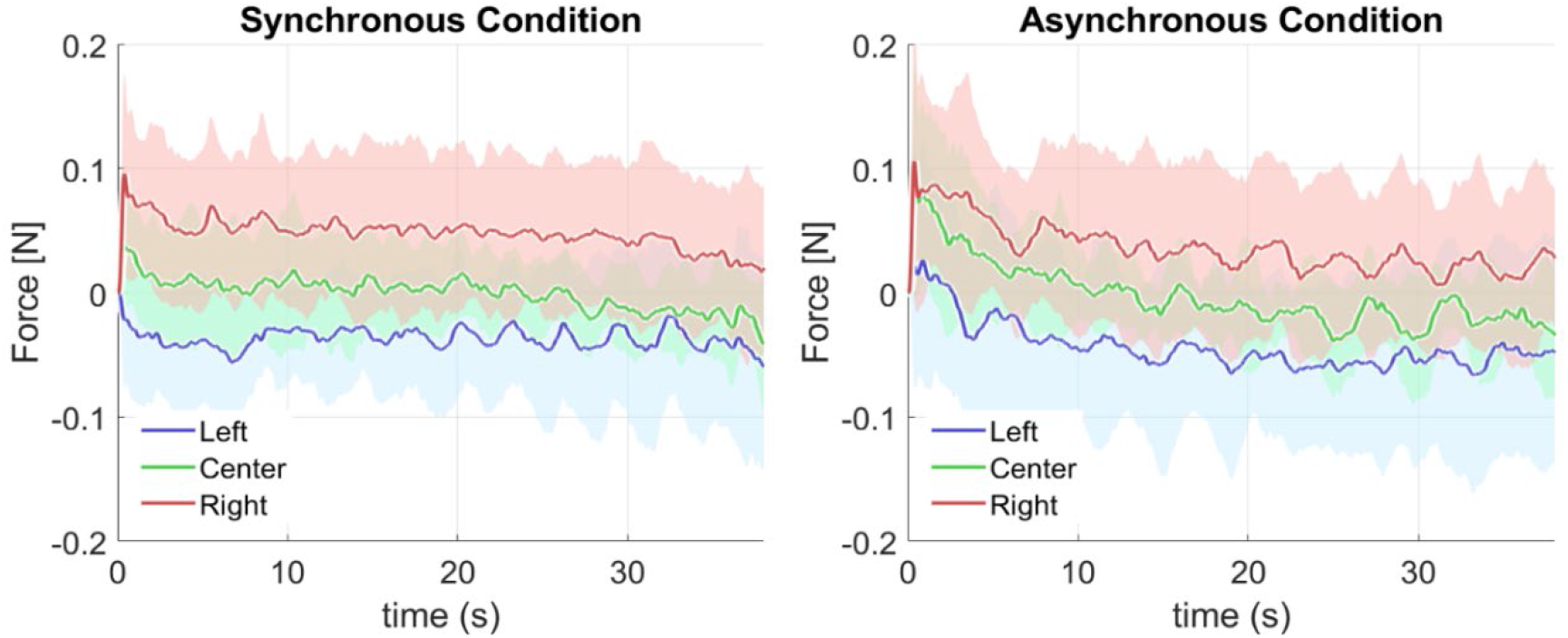
Average participant lateral force exerted during stimulation for the synchronous and asynchronous conditions. The profile is shown separately for the three experimental conditions, Left (blue), Center (green) and Right (red), with solid lines indicating the mean value across participants and the shaded region representing the standard deviation.

Figure 4 shows the registered average force—for the stimulation interval 10s-15s stimulation phase—and standard deviation for all trials and participants for the Left (blue), Center (green) and Right (red) conditions. The marker indicates the mean force and the line expresses the standard deviation. Mean forces in all conditions (Figure 4A-C) showed differentiated directional force patterns towards the virtual hand, in line with the hypothesized trend (purple line in Figure 2A). Figure 4D also shows the mean force for each subject indicating the variability of the forces gains and Figure 4E shows the forces histogram for all subjects split by VR arm location condition. Furthermore, the finding of non-null forces, comparable in intensity, in both left and right directions falsifies the hypothesis of body posture correction or body midline effect.

**Figure 4.**
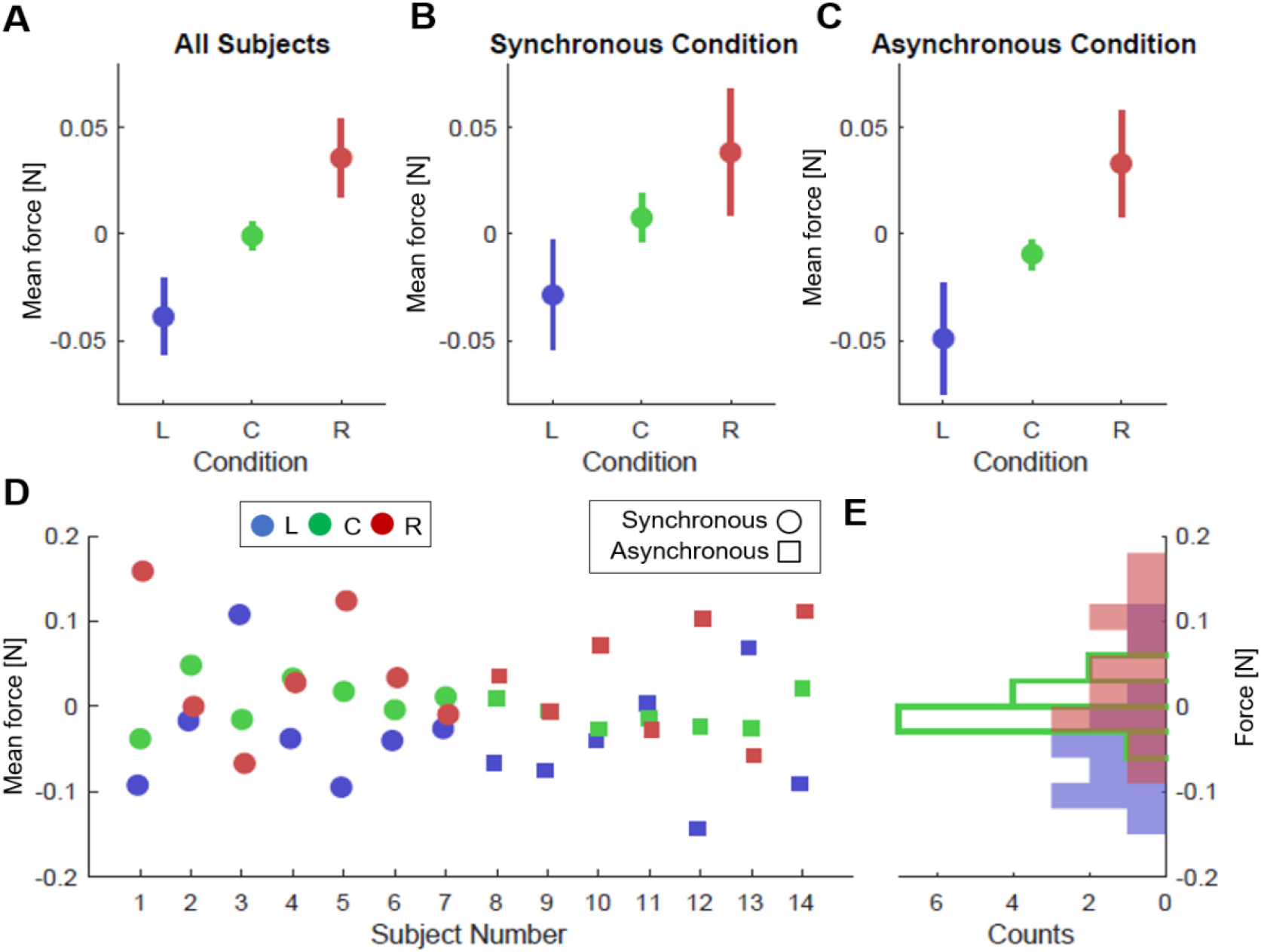
Mean forces applied during stimulation for the different virtual hand location conditions: Left (blue), Center (green), Right (red). The center condition corresponds to the location of the real hand. (**A**) Mean and standard deviation of the force applied by all subjects for the three different virtual hand locations. (**B**) Mean force in the synchronous condition, where tactile (vibrator) and visual stimulation (ball hitting the hand) events were concurrent (i.e. both occurring within less than 100 ms). (**C**) Mean force in the asynchronous condition, where the tactile event was generated randomly along the trajectory of the ball. (**D**) Mean force registered during stimulation for each participant. Three continuous points in the x-axis (L-blue, C-green, R-red) describes one subject. (**E**) Mean force histogram for the three virtual hand location conditions (all subjects).

#### The rubber-hand illusion is active

Our study shows involuntary actions towards the virtual hand during the stimulation phase, despite the inhibitory control and the passive nature of the experiment (participants were instructed not to move). First, we tested the dependence of the applied force, as measured across all trials and participants, on the experimental factors: the virtual hand location, the mode of visuo-tactile stimulation and their interaction. For this, we ran a mixed effect repeated measure ANOVA with virtual hand location (3 levels) as a within-participants factor, VT stimulation modality (synchronous vs asynchronous) as a between-participants factor, and including the possible effect of their integration. We found a significant main effect of the location of the virtual hand (F_2,1674_= 16.05, p=1.2 × 10^−7^). The VT stimulation mode was instead not significant neither as a main factor (F_1,1674_= 1.77, p= 0.18) nor in interaction with the virtual hand’s location (F_2,1674_= 0.18, p= 0.83). We next ran a post-hoc analysis to test our specific hypothesis on the origin of the force dependence on the virtual hand location, i.e., a t-test of the aggregated data across trial repetitions for each participant, taking into account possible inter-individual differences. First, we validated the hypothesis that in the absence of spatial conflict (Center condition) there was no action, i.e., the forces are compatible with a zero mean distribution. A one-sample two-tailed t-test showed that forces in the Center condition have mean 0 (t_1,13_= −1.33, p= 0.90). Second, we tested the hypothesis that in the lateral conditions (Left and Right) forces are different from zero and in the direction of the virtual hand. For this, we ran a one-tailed paired t-tests contrasting forces in each of the later conditions (Right and Left) with those measured in the Center condition. In both cases, our hypothesis is supported by data. For the Right condition the average force is F=0.036 ±0.0009 N, significantly larger (positive) than forces in the Center condition (t_1,13_= 1.84, p= 0.041). For the Left condition the average force is F= − 0.039 ±0.0009 N, significantly smaller (negative) than forces in the Center condition (t_1,13_= −2.00, p= 0.031).

The fact that the ANOVA revealed no effects of the VT stimulation modality indicates that the modulations of the force by the virtual hand location are similar in both synchronous and asynchronous conditions (Fig.4B-C).

### Model predictions vs human observations

Figure 5 shows the comparison between the mean forces registered in the human experiment and the ones predicted by the model. The model was able to predict lateral forces exerted by the participants in the virtual rubber-hand illusion, i.e. for the synchronous group, for the left, center and right conditions. The directions were congruent in all location conditions. In both, the left and the right forces were towards the virtual hand. In the center condition, no lateral forces are present. Statistical differences were only found between location conditions when comparing our model predicted forces with the human data using a two-sample t-test (p<0.05). A mismatch in the strength of the effect was found in the asynchronous condition. While the computational model predicted an attenuated response, in the human experiment similar gains were found in the synchronous and asynchronous conditions. This is also described by the statistical difference in mean of the Left-Asynchronous condition between the model and the human data. This result might be because the VT stimulation adopted in our asynchronous condition was not able to suppress the causal link between the visual input and the body posture inference as intended (see Discussion). To validate this plausible interpretation of our experimental findings we performed additional simulations.

**Figure 5:**
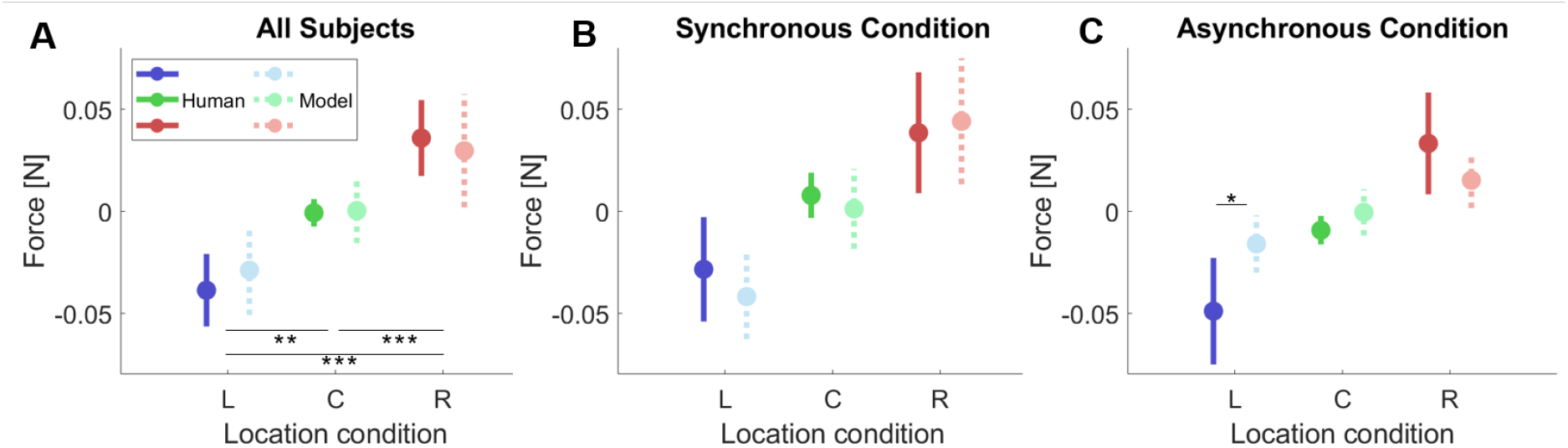
Comparison between the model predictions and human behavior (**A**) Mean force comparison between the human participants and the simulated ones with the active inference model for Left, Center and Right conditions. The continuous line corresponds to the human data and the dashed line with a lighter color to the model predictions. (**B**) Force comparison for the synchronous condition. (**C**) Force comparison for the asynchronous condition.

### Visuo-tactile synchrony evaluation

We further studied the influence of the VT synchrony, taken as a proxy for the level of illusory ownership, on the predicted force. We evaluated the proposed computational model under different degrees of visuo-tactile synchronicity to, first, analyze whether the appearance of forces in the human experiment asynchronous condition may be due to the immersive setting and the formed implicit causal relation between the ball and the hand even using pseudo-random vibration intervals; and second, to assess the influence of the causal parameter *k* in the visual term of the model.

In the classic RHI paradigm, the degree of VT temporal congruency (i.e. the relative timing of events) is known to modulate the illusory experience, and asynchronous conditions are typically adopted to implement control conditions where the illusion is not elicited^9^. However, it has been shown that some degree of asynchrony can be tolerated^12,37^ (e.g. delays up to about 300ms^40^) without breaking the illusion. Furthermore, it has been shown that when experiencing an ownership illusion, the sensitivity threshold for detecting bodily-related VT asynchrony is dampened^41^. Figure 6A shows two sequences of synthetically generated VT events depending on the random strategy used. The visual event refers to the ball on hand and the tactile event to vibration. In practice, in the real experiment, the randomness of the tactile event along the ball trajectory may yield undesired consequences, where the participant assigns causality to both stimuli. This is better captured by the weak asynchronous generation. All together this experimental evidence highlights how the degree of VT asynchrony cannot be mapped one-to-one to a given degree of illusory ownership, which in turn enters in our model via the causal link parameter κ. Thus, to explore the effect of different degrees of illusory ownership on the predicted force, we simulated ten participants varying the causality parameter κ, which depends on the synchrony of the events (see methods) and compared the resulting forces to the ideal asynchronous condition (i.e. the one implying no causal link between visual and tactile stimuli, κ = 0). Hence, a high κ would imply that two events are likely to come from the same source. Figure 6B shows the mean forces and standard deviation of the simulated participants for the asynchronous condition comparing to fixing the causality parameter to 0.3, 0.6 and 0.9.

**Figure 6.**
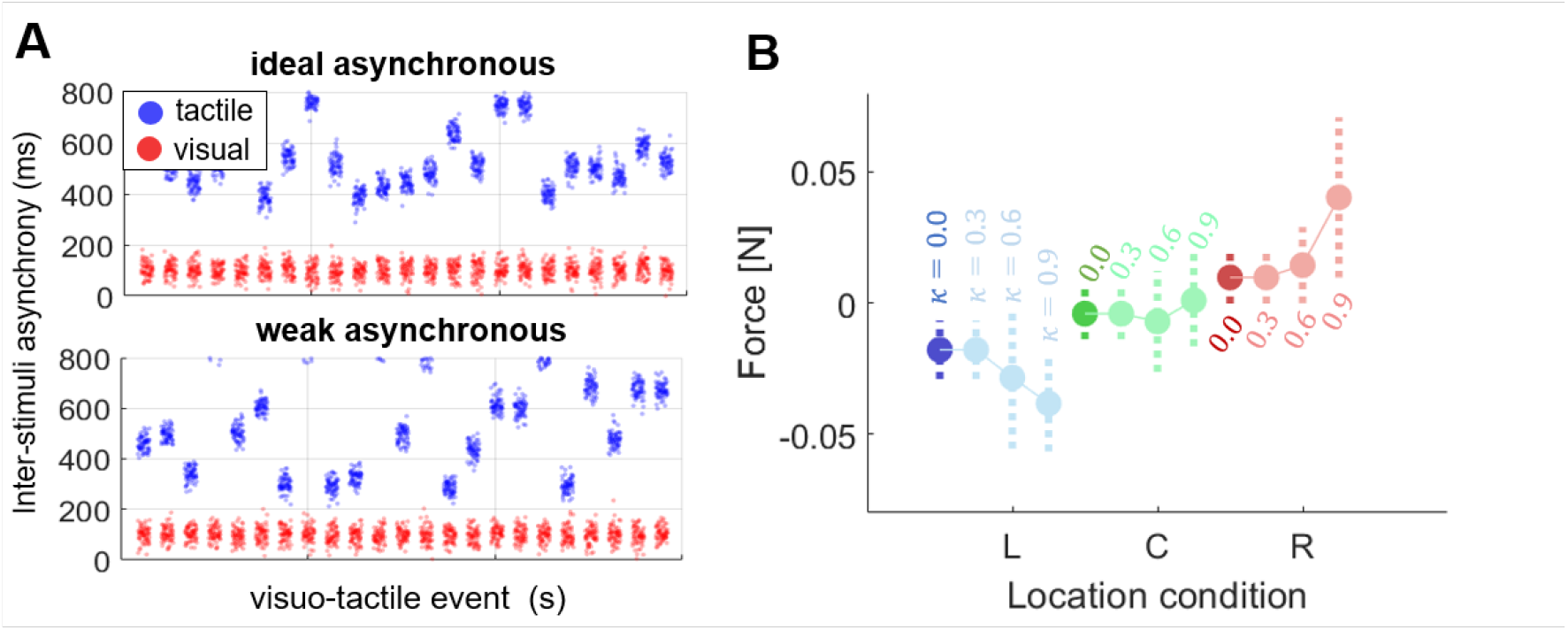
Visuo-tactile stimulation synchrony study. (**A**) Synthetic visual and tactile events in the asynchronous condition depending on the random seed used to generate the vibrations. Y-axis defines how far in time the two events happen. (**B**) Average and standard deviation of the simulated force as obtained for 10 virtual participants each characterized by a different value of the causality parameter κ, which depends on the synchronicity of the visual and tactile events.

Therefore, one of the possible reasons to observe forces in the asynchronous condition is that participants developed an implicit causal relation between the visual virtual arm and their body despite the ball vibrating randomly.

## Discussion

We presented a computational model based on active inference that can predict the active component of the rubber-hand illusion, i.e. a tendency to generate involuntary forces towards the embodied virtual hand, observed in previous experiments and replicated in more details and controlling more factors (under diverse configurations) in the current study. Both model simulations and experimental results provide evidence for an active component of the RHI, which because of its static configuration has been overseen for a long time. This evidence supports the basic interdependence of perception and action.

The present study suggests that humans may move their body to adjust their expected location with respect to other (visual) sensory inputs, which corresponds to reducing the prediction error. This means that the brain adapts to conflicting or uncertain information from the senses by unconsciously acting in the world. We speculate that the proposed theoretical framework can be useful for applications, such as rehabilitation, prosthetics, gaming, etc.

Previous computational models have focused on the perceptual effects of the RHI. For instance, the Bayesian causal model was able to predict the proprioceptive drift by fitting the likelihood to experimental data in humans^42^ and macaques^43^. Our previous predictive coding model^44^ was able not only to replicate the proprioceptive drift but also to show the potential mechanism and the dynamics behind the perceptual effect. The model presented here extends this approach with the inclusion of an active component. Our model, supported by the AIF mathematical framework, can account at the same time for three observed effects in the rubber-hand illusion: proprioceptive^9^ and visual^15^ drifts, and the active component which is the main focus of our study. The perceptual drifts, i.e., proprioceptive and visual, result from a perceptual estimate based on the integration of the two conflicting sensory information, so can be regarded as perceptual biases. The active component or drift (lateral forces) instead may emerge by the active proprioception error minimization through the reflex arc pathway.

Although the free energy principle^26^ (i.e. the basis behind AIF) has been widely proposed to account for several brain processes, such as perception and action^27^, interoception^45^ or self-recognition^46–48^, it is has been difficult to find behavioral validation for its active component. The active inference formulation postulates that both perception and action minimize the surprise, formalized as the discrepancy between our beliefs and the real world. Our model predicts that subjects experiencing body illusions in presence of a visuo-proprioceptive mismatch, tend to generate actions (in the form of exerted forces in the arm) that reduce this discrepancy. During the VHI, the new visual input (virtual hand) is merged with proprioceptive information generating a prediction error on the body location estimation^16^, which can be corrected by an active strategy moving the physical hand to reduce the conflict. The predicted behavior was validated empirically in the human experiment. We observed and measured forces exerted by the physical hand in the direction of the virtual hand. This is, the body may reduce the error of the predicted body location, triggered by the new visual input, by moving the arm. Thus, this could be potential experimental evidence of the active inference framework^49^. However, we should be cautious regarding the model comparison as other computational approaches could also fit our results. For instance, cross-modal learning^50^ could also retrieve these forces by means of sensory reconstruction. Moreover, there are relevant differences between the simplified joint model here adopted and the real musculoskeletal human body.

In contrast with our initial expectations, compensatory forces were observed in our “asynchronous” condition, which was initially expected to be a control condition in which no illusion occurs. The fact that we found a similar active effect (i.e. non-null forces in the direction of the virtual hand) in both synchronous and asynchronous conditions, may be due to our implementation of asynchronous VT stimulation, where the associated sensory conflict might have been not sufficiently high to break the illusion. The accountability of the proposed model to fit human behavioral data depends on the weighting of the visual cue, modulated by the causal parameter *k*, here taken as a proxy for the visuo-proprioceptive binding and indirectly, the level of the experienced illusion. Thus, a limitation of this study is the missing relation between the observed active strategies and the proprioceptive drifts and the strength of the illusion, as we do not explicitly simulate the emergence of the ownership illusion. According to our model, we could expect correlations between the proprioceptive drift and the recorded forces indicating that both effects may be driven by the same underlying process.

Several experiments reported that body ownership is not elicited during asynchronous stimulation^28,51,52^. Such evidence was nevertheless taken from experiments in which other sources of sensory conflict were present (e.g. the spatial mismatch in the classic RHI). In these cases, synchronous visuo-tactile stimulation is required as a trigger for eliciting illusory ownership, while asynchronous stimulation provides additional sensory conflict that keeps the illusion off. Other studies however have shown that VT stimulation is not necessary to trigger the illusion. In fact, ownership illusion can be triggered by congruent visuo-proprioceptive cues only, i.e. for the simple case of spatial overlap between the virtual and real body—as achievable in immersive virtual reality applications—with no further multisensory stimulation^12^. Furthermore, it was shown that once the illusion is established, e.g. through visuo-proprioceptive or visuo-motor triggers, incongruent visuo-tactile stimuli can be tolerated without breaking the illusion^41,53^. This result emerges as a consequence of the fact that the temporal window for VT integration expands due to the causal binding, established by the illusion, between visual stimuli seen on the virtual hand and somatosensory stimuli experienced through the physical hand^41^. It is therefore plausible that the asynchronous visuo-tactile stimulation adopted in our experiment as a control condition did not work as such. This may occur as a consequence of the enhanced immersivity/plausibility of our virtual setup, in combination with repeated exposures to conditions of perfect visuo-proprioceptive overlap between the virtual and the real hands. As reviewed above, previous works show that under this spatial configuration illusory ownership can be sustained even in presence of visuo-tactile asynchronies. Along these lines, it is possible that a residual sense of ownership could be present even in our lateral conditions under visuo-tactile asynchronous stimulation. In this case, the fact that we did not find a significant difference in the forces recorded in the synchronous versus asynchronous conditions, possibly because of a saturation effect whereby even low levels of illusory ownership elicited in the asynchronous condition may trigger perceptual displacements that saturate the active component to the level observed in the synchronous case (in which the exerted forces are very weak).

The interpretation above should be nevertheless taken with cautiousness. Due to experimental requirements (see *Methods*), we did not include explicit assessments of the subjective experience of illusory body ownership. Although lateral configurations in our experiment induced a proprioceptive drift, which is the actual trigger for the predicted force of our model, we cannot properly correlate with the illusion ownership. There is no clear consensus in the literature about this relation. Some works describe a dissociation between ownership and proprioceptive drift based on the lack of a clear correlation between the two^54^. Such lack of correlation appears to be mainly associated with the fact that in visuotactile asynchronous conditions non-null drifts are observed while the reported sense of ownership drops significantly with respect to the synchronous condition. Still, other relevant studies reported clear correlations between subjective ratings of the illusory experience and objective measures of the proprioceptive drift^55–57^. In our pilot study (see *Supplementary Material*) we found that participants exposed to the asynchronous condition, besides displaying a proprioceptive drift, scored positive (although with lower values than in the synchronous condition) to the explicit question on hand ownership (Q3 – “*There were moments in which I felt that the virtual hand was my own hand*”). Similar results with non-negative scores on ownership assessments in visuotactile asynchronous conditions (but significantly lower than in the synchronous condition) have been reported in previous studies^37,41,54^, including in works questioning the causal relationship between ownership and proprioceptive drift^37,54^. Altogether, this might suggest the presence of a residual degree of illusory ownership in our VR asynchronous condition. However, the possibility that the force we measured is dissociated from the sense of ownership cannot be excluded by our experiment alone. In any case, seen in the larger framework of the experimental evidence on ownership illusion from the last decade, our results provide new evidence for the active component of the RHI and novel insights for understanding its underlying computational base.

Previous work has shown the strong link between self-perception and action. This coupling has been observed in reactive responses to the threat observed on an embodied fake body^11,58^. Further support comes from experimental evidence showing how the sense of agency over the movements of virtual hands or full body avatars enhances the sense of illusory body ownership^53,59,60^. Our results point to a possible complementary effect, i.e. to how the sense of body ownership can potentially affect motor behavior, depending on the sensory cues available. Further studies in which the subjective sense of ownership is explicitly assessed are nevertheless needed to ascertain this possibility.

Our model provides an active inference perspective on motor behavior showing how the need to suppress sensory conflict could trigger movement, as an active strategy to preserve the integrity of self-perception. We expect that such sensory-conflict-driven actions could affect motor performance in goal-directed, as in the case of the self-avatar follower effect observed in VR settings^19^. Our proposed perception-action coupled approach would be furthermore advantageous for modeling adaptation during interaction, as world/body variability can be instantly taken into account while maintaining a coherent body pose estimation.

Future research needs to address the computational model behind the dynamics of illusory experience itself and more realistic visual input^61^. One possibility could be to integrate, in a hierarchical fashion, our AIF model with Bayesian causal inference models previously proposed for body ownership illusions^8,42^. These models are based on the assumption that the illusory sense of body ownership arises when the brain attributes the visual information from the rubber or virtual hand to the same entity that gives origin to somatosensory input. The causal inference model we propose here, instead, simulates the perceptual process inferring the location of the hand and the origin of actions that minimize sensory conflicts while assuming that the agent believes to some degree (represented by the *k*parameter) that the visual and the proprioceptive/tactile sensory input are associated to the same origin. The observation of forces in the asynchronous condition in human experiments reflects the need for further testing the model under several control conditions and for integrating the two models. Even more importantly, observing the active component in a non-static environment will be an important step to corroborate the robustness of the proposed model. Heading in this direction, we plan to test the model in conditions where the real hand is able to move, and further when the virtual arm can be controlled by the agent.

## Methods

### Computational model

We model body perception as inferring the body posture by means of the sensation prediction errors (visual and proprioceptive) and the error in the predicted dynamics. Inspired by the fact that the RHI affects the joint angles perception^62^, body posture is defined by the joint angles. To design the arm model we only consider one degree of freedom: the elbow that rotates over the vertical axis (Figure 2A). Thus, we define *s*_*p*_ as the measured/observed joint angle of the elbow and *μ*^[0]^ ∈ (− π/2, π/2) as the inferred elbow joint angle. The subscript notation reflects the order of the dynamics. *μ*^[0]^, *μ*^[1]^, *μ*^[2]^ represents the position, velocity and acceleration of the inferred brain variables^63,64^. Visual information is given by the horizontal location of the center of the virtual hand *s*_*v*_.. We further define that zero degrees measurement indicates when the arm is in perpendicular to the horizontal axis and parallel to the body sagittal plane. Hence, left and right rotations are negative and positive respectively. Given the length of the arm *L*, the generative model that predicts the hand visual location depending on the joint angle (*μ*^[0]^) is as follows:

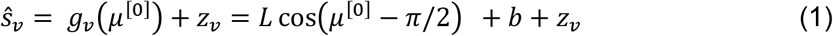

where *b* is a localization perceptual bias (sampled from human data^15^) that depends on each participant and *z*_*v*_ is normally distributed noise. The rest of the observations can be predicted with a Gaussian with mean 0. Thus, the observed sensations ***s*** and the generative model *g*(*μ, v*) that predicts the sensations are:

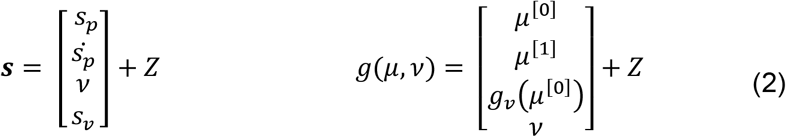

Where *s*_*p*_ and 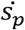 are the joint angle and velocity, *v* is the causal variable, i.e., the perceived location of the virtual hand, *s*_*v*_ is the measured position of the virtual hand and *Z* is the noise associated with each variable. Note that during the RHI experiment the participant does not have visual access to their hand.

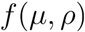

does not expect any action as there is no task involved and it is modeled as a mass-spring system. Note that this function is an approximation of the real model of the arm:

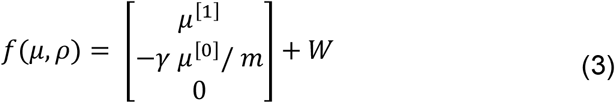

Where *W* is the associated noise of the process, *m* is the mass of the arm and *γ* is the viscosity parameter. In the case of having a task we shall include a perceptual attractor in the sensory manifold and its transformation to the joint variables. For instance, we can include the virtual arm as the goal by substituting in equation (3) the second row by (*Tμ*^[0]^)*A*(*μ*^[0]^, *ρ*) − (*γμ*^[0]^)/*m*, where is *A*(*μ*^[0]^, *ρ* = *β* (*ρ* − *g*_*v*_(*μ*^[0]^)) and the *T*(*μ*^[0]^ = −*L* sin (*μ*^[0]^ − *π*/2). However, in this case we do not have a desired goal.

We infer the elbow angle by optimizing the free-energy bound under the Laplace approximation^65^. Defining the error between the inferred brain variables and the dynamics generative model as ***e***_*μ*_ = ***μ*** − *f*(*μ, ρ*), the differential equation that drives ***μ*** = [*μ*^[0]^, *μ*^[1]^, *μ*^[2]^]^*T*^ is:

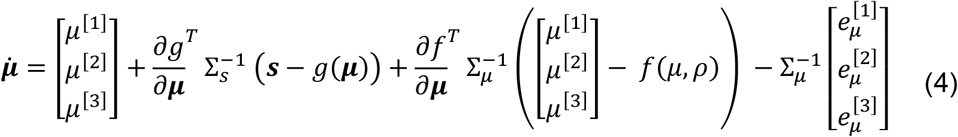

Regarding the action, it only depends on the sensory input, as during the RHI the participant only relies on the proprioceptive input. Thus, we define its differential equation as:

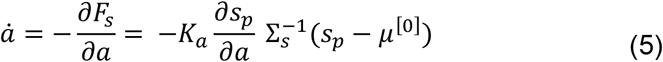

Where *K*_*a*_ is the gain that modulates the step size of the differential equation (similar to gradient descent). Note that although we computed the action and use it as an estimate of the force applied at the level of the arm, we did not use it in the update of the real hand position, to simulate the constraint of the RHI where the real hand is fixed and cannot move.

Experiments have shown that during the RHI participants modify the precision of visual and proprioceptive cues^14^. Thus, we optimize their precision also to reduce the prediction error:

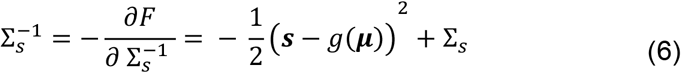

We set a minimum of exp(1) and exp(0.1) for visual and proprioception precision respectively.

Finally, we considered that the perception of the virtual hand location is another unobserved variable. Thus, the model also infers the visual horizontal location, allowing the blending of the real hand perception error with the virtual hand and therefore, producing a visual drift.

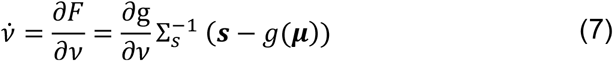

To generate individual differences between participants we fixed all parameters and we randomly selected the initial proprioceptive variance 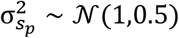. This means that each participant has a different precision of the proprioceptive cue. We further included the bias in the predicted real hand location based on reported data of RHI human experimentation (in cm) *b* ∼ 𝒩(0,4.2794). The length (in cm) of the forearm was drawn from a normal distribution *L* ∼ 𝒩(44.7208,2.4807). Bias and the forearm lengths data were taken from participants’ real data^15^. During stimulation, we also introduced artificial noise in s_p_ and s_r_ readings.

Finally, to generate the effect of synchronous and asynchronous stimulations on the illusion and therefore on the predicted force, we introduced the parameter *k* as the probability of touch and visual cues being generated by the same source, so as an estimate for the level of illusory ownership over the virtual hand^8,42^ The *k* parameter weights the error between the expected visual location of the arm and the virtual hand location input: *k*(*v* − *g*_*v*_ (*μ*)). The synchronous and asynchronous VT stimulation were synthetically implemented producing tactile and visual events in the ranges of 100 ms in the synchronous condition and 800 ms otherwise^34^, and we considered these as two extreme cases where the illusion was either saturated or completely suppressed. So a *k* value of 1 was set for the synchronous case, meaning that the visual input is generated by our body, and a *k* value of 0.3 was set for the asynchronous condition, assuming that in this case the virtual hand was perceived as an external object not related to the real body. In addition we performed additional simulations, adopting different values of *k* to explore how different degree of the illusion intensity would affect the forces applied at the level of the real arm.

### Devices

Subjects were seated with their shoulders restrained against the back of a chair by a shoulder harness and their hand and forearm were rested by a flat surface attached to the vBOT robotic manipulandum^66^ with their forearm supported against gravity with an air sled (Top view Figure *7*; Lateral view Figure 1C). The robotic was locked to prevent movements of the participants’ hand while measuring the applied force. Position and force data were sampled at 1KHz. Endpoint forces at the handle were measured using an ATI Nano 25 6-axis force-torque transducer (ATI Industrial Automation, NC, USA). The position of the vBOT handle was calculated from joint-position sensors (58SA; IED) on the motor axes. Visual feedback was provided using a virtual reality (VR) device (Oculus Rift V1, Facebook technology). When using the VR system, any visual information about the real body location was prevented. The virtual environment was designed and programmed in C# using the Unity engine. The virtual environment and body size were fixed for all participants (e.g. the length of the VR arm was always the same). Tactile feedback was provided with commercially available coin vibrators (model 10B27.3018) placed on the middle point of the participant’s hand dorsum, controlled with an embedded chip and connected via USB to the VR engine. The timing of the vibration events is controlled in relation to a visual animation of a virtual ball bouncing on the dorsum of the virtual hand. A random number was generated every time the ball contacted the virtual arm. This random number was translated into the timing to trigger the tactile event along the trajectory of the ball.

**Figure 7:**
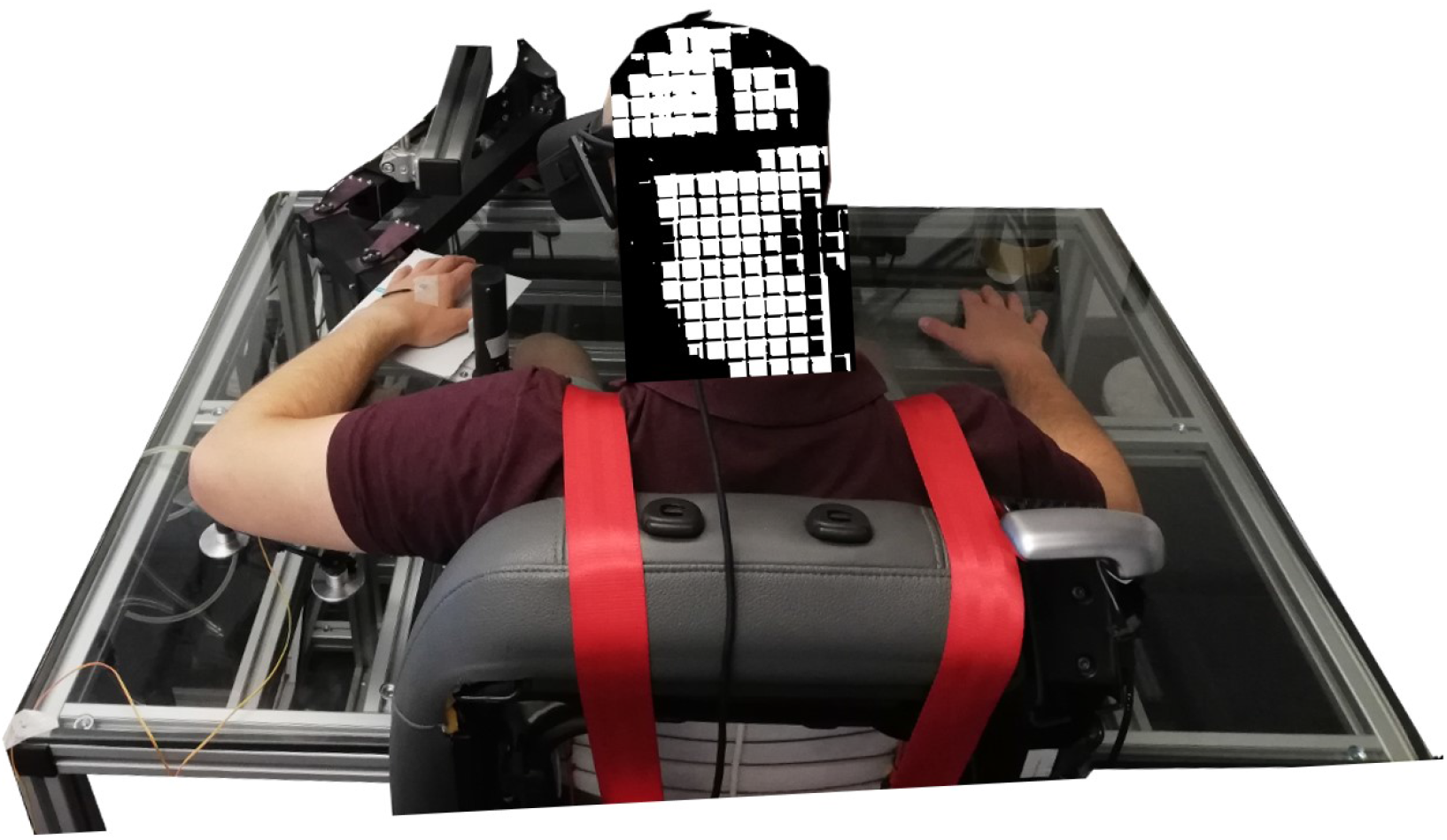
Experimental setup. Back-top view of a participant on the vBOT manipulandum with the VR system and the coin vibrator attached to the dorsum of the left hand.

### Ethics

Informed consent was obtained from all participants, and the study was approved by the ethics committee of the Medical Faculty of the Technical University of Munich, and adhered to the Declaration of Helsinki. The study was performed in accordance with the ethical guidelines of the German Psychological Society (DGPs).

### Participants

Fourteen right-handed volunteers (seven male and seven female) from 18 - 40 years old with normal or corrected-to-normal vision participated in this study. Handedness was evaluated using the Edinburgh Handedness Inventory. All participants had no neurological or psychological disorders, virtual sickness, nor skin hyper sensibility as indicated by self-report A pretest with the vibrator was performed on the participant hand who gave a written statement that the vibrator did not harm himself. We ensured that all participants were not wearing nail polish or had remarkable visual features on the left or right forearm and hand. Participants’ height was between 1.6 and 1.9 m to fit within the constraints of the experimental apparatus. Participants were economically compensated with eight euros per hour.

### Experimental design and procedure

First, we randomly assigned seven participants for the synchronous and seven for the asynchronous condition. In total, we registered 1680 trials or samples across all participants. The generation of the conditions was computed randomly in blocks as described in Figure 8.

**Figure 8:**
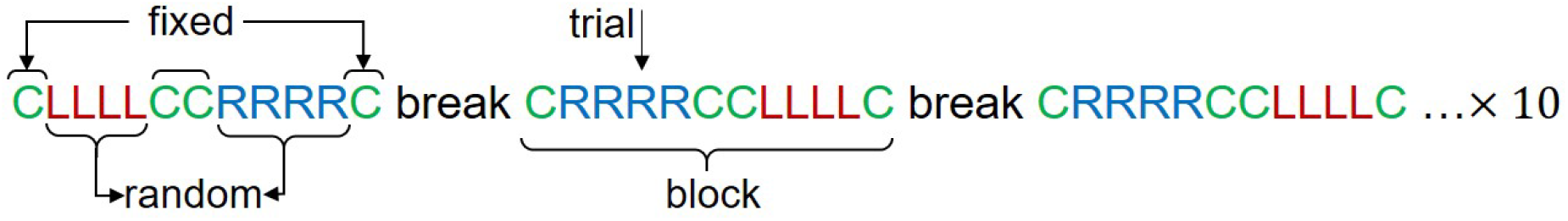
Generation of block and condition order for participants. Each participant did 10 blocks of 12 trials (4 for each VR location condition). Center trial positions were fixed while the right and the left set of four trials were randomly distributed between the center trials.

The center condition was always fixed in the beginning, the middle and the end of every trial. Four trials of the left or right condition were randomly selected between the center conditions. The selection of the center condition in the middle was imposed to reduce the localization bias (i.e. the effect of proprioceptive drift induced in the previous trial on the following ones) produced when stimulating in the left and right conditions.

Each trial corresponded to 40-seconds of VT stimulation where the force was recorded. To register the samples, we performed 10 blocks per participant separated by a short break. Each block was composed of 12 trials (4 trials for each virtual hand location condition, left, center, right). In total, each participant performed 120 trials (40 per visual hand location condition).

Participants were seated on the experiment chair, in front of a table as described in Figure 7, following the instructions of the experimenter. Once the chair was adjusted to the needed height, the experimenter attached the vibrator to the dorsum of the hand. Participants then wore the Oculus Rift VR system. The left hand rested on the vBOT manipulandum and the right hand on the table (Figure 7). Participants were asked to stay relaxed and still in the chair.

The initial location of the left hand was fixed for all participants and was programmed in the Manipulandum. Each trial consisted of two phases and then a resting period:

1. *Cross*: At the beginning of each trial participants were asked to look frontal towards a cross that appeared in a random location around the middle of the scene to avoid the use of the head angle as a prior cue.
2. *Stimulation*: While participants were looking at the cross, a virtual arm in the Left, Center or Right condition appears in the virtual scene with a ball on the top of the hand. After this appears in the virtual scene, participants were asked to look down at the index finger of the left hand. Participants can only see the virtual hand once they look down. In the synchronous condition, every time the ball touches the virtual hand there is a tactile stimulus through the vibrator (a touch event every two seconds approximately). In asynchronous stimulation, the vibration occurs with a random delay after the visual stimulation.
3. *Resting*: At the end of each block, participants rested for 1 minute alternating removing the VR system and moving the arms without removing the VR.

We tested both synchronous and asynchronous VT stimulation. In the synchronous condition, the events of ball contact with the virtual hand and the vibration on the real hand occurred within the temporal window for perceived simultaneity (less than 100 ms difference). On the other hand, the asynchronous condition vibration events were generated randomly between the visual hit and the highest height reached by the ball.

Data were analyzed offline using Matlab (R2019a) for data cleaning and preprocessing and using R for statistical testing. For force measurements we performed offset removal by subtracting the mean force over each 9-minute block of trials (containing equal numbers of all three condition types) from the force data, to ensure that a bias on the force values was not introduced through shifts in participant limb posture. The impact and interaction of experimental factors on the observed forces was assessed running a mixed effect repeated measure ANOVA, while paired t-tests were run for specific post-hoc hypothesis testing.

### Code and data

For reproducibility of the results, we provide an instance of the developed model (in python, Google colab) with fixed parameters that can be executed in the following link and a Jupyter notebook in the supplementary material. Force data that support the findings are provided in the supplementary material. Manipulandum raw data is available from the corresponding author upon request.

## Supplementary information

For completeness, we include supplementary experimental results of a preliminary study to assess the proprioceptive drift and level of body-ownership induced with our experimental setup. See supplementary material.

## Acknowledgments

The authors would like to thank Gordon Cheng for his advice and support, and Mohamad Atayi for the development of the VR environment and his help in the preliminary study. This work was partially supported by the MSCA SELFCEPTION project (www.selfception.eu) EU H2020 grant no. 741941.

## Contributions

P.L. conceived of the idea, developed the computational model and performed the model analysis. P.L., S.F., D.F. designed and planned the experiments. P.L. and S.F. collected the data. A.M. and D.F. performed the human analysis. P.L. and A.M. drafted the manuscript. P.L and D.F. designed the figures. All authors revised the manuscript.

## Supplementary information

### Body-ownership and proprioceptive drift study

We performed a preliminary study to assess the proprioceptive drift and level of body-ownership induced with our experimental setup. Eight participants (a different group from the main study) were tested under the same experimental setup as for the action paradigm. We measured the proprioceptive drift and the body-ownership score. The experiment had ten blocks (five synchronous and five asynchronous), where the virtual hand was placed in one of the three locations: Left, Center, Right. The implementation of the synchronous and asynchronous conditions was the same as in the main experiment.

Equally as the main experiment, participants were seated on the experimental chair, in front of a table, following the instructions of the experimenter. Once the chair was adjusted to the needed height, the experimenter attached the vibrator to the middle point of the hand dorsum. Participants then wore the VR system. The left hand rested on the air sled and the right hand rested on the table. The initial location of the manipulandum was fixed and programmed for all participants. Each condition trial consisted of four phases: 1) *Pre-localization*: Only a table was displayed in the VR and a vertical ruler (depending on the condition) appeared. The participants had to indicate verbally the number on the ruler where they currently perceived the location of the index finger of their left hand. This process was repeated 8 times. As in the main force experiment, between each measurement, participants were asked to look frontal at a cross that appears in the middle of the scene. To avoid the use of the head angle as a cue we injected random noise in the cross location. 2) *Stimulation*: While looking at the cross the virtual arm appeared in the scene in the correct location condition with a bouncing ball on the top of the hand; Participants were instructed to look down towards their index finger. 3) *Post-localization*: Participants were asked to indicate (say the number) where they perceived the index finger of the hand in the same horizontal or vertical ruler as stage 1; Finally, 4) *Resting*: at the end of each trial, participants removed the VR system and rested for 1 minute while filling the illusion questionnaire.

#### Proprioceptive drift and body-ownership

As expected, proprioceptive drifts were found in the left and right conditions (Figure 1A). Perceptual drifts effects were also found in the asynchronous condition indicating that random vibrations were also integrated although to a lesser degree. The virtual immersion made that even in the asynchronous condition participants experienced some degree of partial body-ownership, as emerging from by the subjective evaluation of their level of body-ownership towards the virtual hand. Figure 1B shows the questionnaire results using a seven-point Likert scale (3 indicating strong agreement and –3 indicating strong disagreement). Although participants could detect the temporal misalignment of the VT stimulation in the asynchronous condition, as shown by scores to question Q2, this did not completely prevent participants to experience ownership illusion over the virtual hand also in the asynchronous condition (as shown in the positive values of Q1 and Q3).

#### Questionnaire

The illusion questionnaire was developed by adapting the one presented in Nina et al.^1^ to virtual environments with previously developed questionnaires for virtual reality RHI^2,3^. It is compounded by nine questions, where Q1-Q3 describe the RHI effect^4^ and Q4-Q9 are control questions.

1. It seemed as if I was feeling the vibration in the location of the virtual arm.
2. Sometimes I had the sensation that the vibration I felt in my hand was caused by the contact of the ball with the virtual hand.
3. There were moments in which I felt that the virtual hand was my own hand
4. There were moments where the touch I was feeling came from somewhere between my own hand and the virtual hand.
5. There were moments in which I felt as if my real hand was becoming virtual
6. It seemed as if I might have more than one left hand
7. The virtual hand started to look like my hand, in terms of shape, skin tone, freckles or some other visual aspects.
8. I felt as if the virtual hand was drifting towards the real hand.
9. I felt as if my real hand was drifting towards the virtual hand.

